# Wing damage affects flight kinematics but not flower tracking performance in hummingbird hawkmoths

**DOI:** 10.1101/2020.08.16.252759

**Authors:** Klara Kihlström, Brett Aiello, Eric J. Warrant, Simon Sponberg, Anna Stöckl

## Abstract

The integrity of their wings is crucial to the many insect species that spend distinct portions of their life in flight. How insects cope with the consequences of wing damage is therefore a central question when studying how robust flight performance is possible with such fragile chitinous wings. It has been shown in a variety of insect species that the loss in lift-force production resulting from wing damage is generally compensated by an increase in wing beat frequency rather than amplitude. The consequences of wing damage for flight performance, however, are less well understood, and vary considerably between species and behavioural tasks. One hypothesis reconciling the varying results is that wing damage might affect fast flight manoeuvres with high acceleration, but not slower ones. To test this hypothesis, we investigated the effect of wing damage on the manoeuvrability of hummingbird hawkmoths (*Macroglossum stellatarum*) tracking a motorised flower. This assay allowed us to sample a range of movements at different temporal frequencies, and thus assess whether wing damage affected faster or slower flight manoeuvres. We show that hummingbird hawkmoths compensate for the loss in lift force mainly by increasing wing beat amplitude, yet with a significant contribution of wing beat frequency. We did not observe any effects of wing damage on flight manoeuvrability at either high or low temporal frequencies.

## Introduction

Insects are masters of flight – with their fragile chitinous wings they perform impressive aerobatic manoeuvres, such as a dragonfly catching its prey in the air, or a hawkmoth hover-feeding from a flower which is moving in the wind. Yet, wing wear caused by collisions or predation is almost unavoidable for most insect species. The consequences of wing damage on flight kinematics have been investigated in variety of insect species – with disparate results depending on species and the type of damage inflicted. A general trend across these species is that reduced wing area, and consequently lift force (Ellington 1984a), is compensated for by increasing wing beat frequency (bumblebees: (Hedenström et al. 2001, Haas & Cartar 2008), hawkmoths: (Fernández et al. 2012, Fernández et al. 2017) and butterflies (Kingsolver 1999)). Force production also scales with increased wing beat amplitude, which might therefore also compensate the loss in lift force (Ellington 1984a). However, wing beat amplitude does not change in bumblebees with either symmetric or asymmetric wing damage (Hedenström et al. 2001), and only upon asymmetric wing damage in the hawkmoth *Manduca sexta* (Fernández et al. 2012, Fernández et al. 2017).

While the changes in wing beat kinematics upon wing damage are rather similar across insect species, their effects vary considerably. In hawkmoths, both symmetric and asymmetric wing damage increases the metabolic cost of hovering (Fernández et al. 2017), while this is not the case in bumblebees (Hedenström et al. 2001). However, bumblebees with damaged wings have been shown to have a higher mortality (Cartar 1992). It was therefore hypothesised that wing damage might impair flight performance rather than the energetic balance, thereby affecting fitness. However, no changes in foraging performance, nor flight speed, acceleration, and distance to the ground, were observed upon artificially inflicted wing damage in a simple foraging task (Haas & Cartar 2008). Moreover, only the animal’s peak acceleration, but not their ability to avoid obstacles, was affected in a more complex foraging task (Mountcastle et al. 2016). Similar to bees, no fitness consequences were observed in foraging butterflies (Kingsolver 1999), while a clear decrease in vertical acceleration and average velocity and prey capture success could be documented in wing-damaged dragonflies, which require complex and fast aerial manoeuvres for prey capture (Combes et al. 2010). Thus, one might hypothesise that wing damage affects fast aerial manoeuvres more than steady flight performance or take-off and landing.

We therefore decided to investigate the effect of wing damage on the manoeuvrability of a hawkmoth. Even though their hovering kinematics (including energy requirements with intact and damaged wings) have been well studied, little is known concerning how wing damage affects flight performance. Since hawkmoths forage on the wing and extract nectar while hovering in front of flowers, they need to track wind-induced flower movements using fast corrective flight manoeuvres (Farina et al. 1994, Sprayberry & Daniel 2007, Sponberg et al. 2015, Stöckl et al. 2017). Hawkmoths are an ideal study system for studying the flight performance consequences of wing damage. We chose to study the hummingbird hawkmoth, *Macroglossum stellatarum*, a species whose individuals hibernate as adults and therefore have lifespans of several months (Pittaway 1993), with the consequential risk of wing damage during this time. To investigate the effects of wing damage on their manoeuvrability, we tracked hawkmoths feeding from an artificial flower that could be moved at specific temporal frequencies (Fig. 1A). We hypothesised that wing damage is compensated in hummingbird hawkmoths by adjustments in wing beat frequency and amplitude, similar to other insects. Moreover, we expected to find effects of wing damage on flower tracking performance, in particular at the higher temporal frequencies that require quicker turning manoeuvres and correspondingly higher peak accelerations, in line with similar results in dragonflies and bumblebees.

**Fig. 1.**
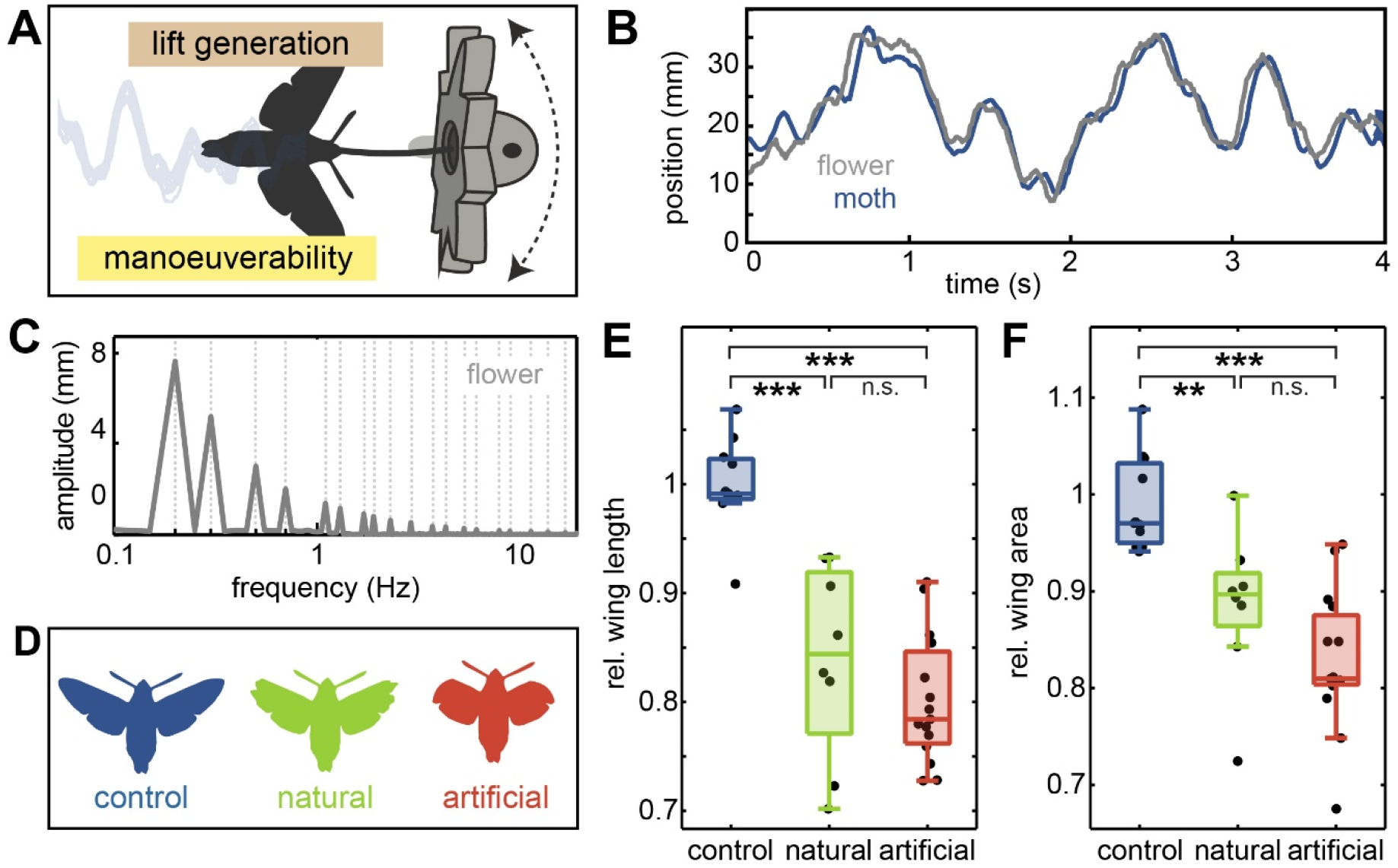
The role of wing damage on hawkmoth flight performance. **A** Wing shape and wing area could affect a hawkmoth’s foraging performance both by impairing the lift force generation required to keep the animal airborne, as well as by impairing the manoeuvrability of the animal as it tracks the movement of the flower. **B** To test the manoeuvrability of the animals, we moved a robotic flower at specific temporal frequencies while the animals were hover-feeding from the flower and attempted to track it. **C** The amplitude of flower movements was adjusted to keep the velocity at the different frequencies constant. **D** To study the effect of wing damage on flight and flower tracking performance, we compared a control group with intact wings to a group with natural wing damage, and a group in which we clipped the distal forewings to the maximum extent possible that allowed the animals to fly. **E-F** Forewing length and forewing area were normalised by the expected forewing length and area given each animal’s body length (see Methods, Fig. 1 – S2). Black dots denote individual hawkmoths, boxplots depict the median and 25^th^ and 75^th^ percentiles of the samples. Whiskers represent the data range excluding outliers (values extending 1.5 interquartile ranges beyond the upper and lower box limits). Statistical differences between groups are indicated as: *** p < 0.001, ** p < 0.01, * p < 0.05, n.s. p > 0.05 (ANOVA with Tukey’s HSD corrected post-hoc test was performed after testing the linearity of residuals, Table S2).

## Methods

### Animals

Wild adult *Macroglossum stellatarum* L. (Sphingidae), were caught in Sorède, France. Eggs were collected and the caterpillars raised on their native host plant *Gallium sp*.. The eclosed adults were allowed to fly and feed from artificial flowers similar to the experimental flowers, in flight cages (70 cm length, 60 cm width, 50 cm height) in a 14:10 h light:dark cycle for at least one day before experiments.

### Experimental groups

Three different experimental groups were tested: *control* animals with intact wings, animals with *natural* wing damage which was caused by flying in their holding cages, and *artificial* wing damage (Fig. 1D) induced by cutting the distal tip of the forewing of the hawkmoths, following the shape of their hindwing for consistency to reach an average reduction of 19% in wing area and 25% in forewing length – the maximum amount of wing damage that would still allow the hawkmoths to take off (Table 1). Animals were allowed to recover for one day in their holding cages before experiments started.

We photographed every animal after the experiment, and determined their body and wing size using Fiji software (Schindelin et al. 2012). Their total body length was measured along their anterior to posterior extent, the total wing length was measured for both wings, from the wing joint to the tip of their forewing, and the area of both wings was quantified by tracing with the polygon tool in Fiji. The data from the left and right wing was averaged for further analysis.

When reporting wing damage, we accounted for the individual size of the animals – and thus the individual size of their intact wings - by normalising the measured wing morphology of the three treatment groups with respect to the intact wing morphology expected for an animal of this size. This was possible because wing length and wing area scaled tightly with animal length (Fig. 1 –S2). We therefore computed the allometric scaling between wing length (Fig. 1 –S2A) and wing area (Fig. 1 – S2B) using animals with intact wings (the control animals in this study, as well as an additional 20 animals with intact wings that were not tested with the robotic flower). To obtain the scaling exponent *b* and the scaling constant *a* of the allometric relationship

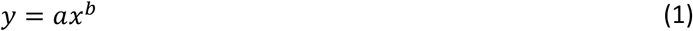

we used Model II (reduced major axis) regression implemented in the *gmregress* script for MATLAB (A. Trujillo-Ortiz, https://www.mathworks.com/matlabcentral/fileexchange/27918-gmregress, retrieved March 19, 2020) to fit the parameters in the log-transformed version of the equation:

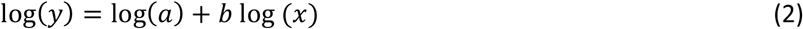

This yields the scaling exponent *b* as the slope of the linear relationship, and the log-transformed scaling constant *a* as the y-axis intercept (Warton et al. 2006). With b=0.831 and a=1.397 for wing length, and b=1.831 and a=0.262 for wing area, we could use animal size to calculate the wing length and area that would be expected for the individuals in all treatment groups given their body length. These expected values were then used to normalise the measured wing length and area (Fig. 1E,F), so they represent the proportion of expected wing length and area for an animal of a given body length.

### Experimental setup

We used a robotic flower assay as our experimental setup. This assay was pioneered by (Farina et al. 1994, Sponberg et al. 2015), and also used in (Roth et al. 2016, Stöckl et al. 2017, Dahake et al. 2018). A flight cage of the same size as the holding cage was lined with soft muslin cloth and covered with black cloth on the three sides, while the front and top were sealed with Perspex plates for filming. A 3D-printed plastic flower (48 mm in diameter, on a 140 mm stalk) was placed at the centre of the flight cage. Placed at its centre was a nectary with an 8.3 mm opening, which was filled with 10% sucrose solution. The flower could be moved sideways in shallow arcs around the central pole, such that the primary motion was a lateral translation. The movement was controlled by a stepper motor (0.9 degree/step resolution, 1/16 microstepping, Phidgets, Inc.) and a custom-written Matlab program. The cage was illuminated from above with an adjustable white LED panel and diffuser (CN-126 LED video light, Neewer). The light intensity was set to 3000 lux (measured with a ScreenMaster, B. Hagner AB, Solna, Sweden, at the position of the artificial flower). In addition, two 850 nm IR LED lights (LEDLB-16-IR-F, Larson Electronics) provided illumination for the infrared-sensitive high-speed video cameras (MotionBLITZ EoSens mini, Mikrotron), which was used to film the experiment. Videos were recorded at 100 fps, allowing us to record sequences of up to 28 seconds to analyse flower tracking.

### Behavioural experiments

Individual moths were taken from their holding cage and introduced into the experimental cage. Here they were given 5 minutes to warm up their flight muscles and take flight. Most hawkmoths would approach the artificial flower within a few minutes after taking off. When their proboscis contacted the nectary, we started moving the artificial flower. If the animals did not take flight, or did not feed from the flower within 10 minutes of taking off, we aborted the experiment and tested them again the next day. To move the artificial flower, we used a “sum-of-sines” stimulus of 20 s duration comprising a pseudo-random sum-of-sine stimulus composed of 20 temporal frequencies, which were prime multiples of each other to avoid harmonic overlap: 0.2, 0.3, 0.5, 0.7, 1.1, 1.3, 1.7, 1.9, 2.3, 2.9, 3.7, 4.3, 5.3, 6.1, 7.9, 8.9, 11.3, 13.7, 16.7, 19.9 Hz. High frequencies had lower amplitudes and vice-versa, to assure equal peak velocities at all frequencies and avoid saturation due to power limitations (Roth et al. 2014) (Fig.1C).

### Flower tracking analysis

The positions of the flower and the hawkmoth were digitised from the videos using the DLTdv5 software for Matlab (Hedrick 2008) as described in (Sponberg et al. 2015, Stöckl et al. 2017). In the top view, a point on the flower and a reliably identifiable point on the thorax of the moth were used for reference. To analyse changes in body pitch angle, we also tracked the tip of the abdomen for the first 200 frames of the stimulus. In the dorsal camera view, an increase in body pitch angle should manifest as a decrease in the distance between the thorax and abdomen tip (Fig. 2-S2). We only analysed sequences where the hawkmoth’s proboscis was in contact with the nectary.

**Fig. 2.**
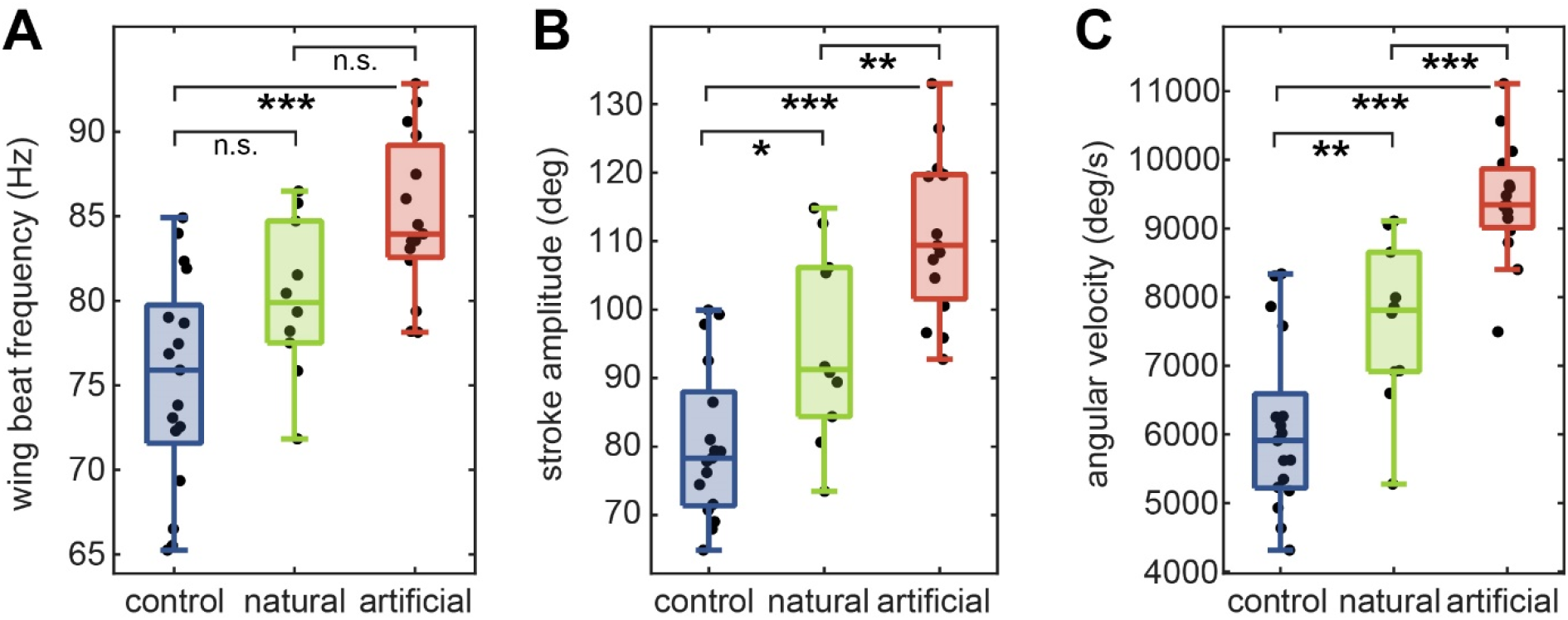
Wing beat parameters of the three treatment groups. **A** Wing beat frequency and **B** horizontal projection of wing stroke amplitude measured during a hovering at the flower stimulus. **C** Wing stroke velocity was calculated as the product of wing beat frequency and amplitude. Statistical differences between groups are indicated as: *** p < 0.001, ** p < 0.01, * p < 0.05, n.s. p > 0.05 (ANOVA with Tukey’s HSD corrected post-hoc test was performed after confirming linearity of residuals, Table S3).

To analyse the tracking performance across the entire stimulus, we extracted the absolute Euclidian distance between the hawkmoth’s thorax and the nectary of the flower to obtain a measure for the average displacement of hawkmoth and nectary (Fig. 5A). Moreover, we calculated the absolute length of the path the hawkmoth’s thorax travelled, relative to the path length of the flower (Fig. 5B). To analyse the tracking performance at each temporal frequency of the stimulus, we extracted the amplitude and phase components in the corresponding power spectra at the stimulus frequencies. As in previous studies with the same stimulus (Sponberg et al. 2015, Stöckl et al. 2017), we did not analyse data at the highest two temporal frequencies (16.7, 19.9 Hz), moths did not reliably track these frequencies. We used a system identification analysis (Cowan et al. 2014) to characterise the flower tracking performance of the hawkmoths as described previously (Sponberg et al. 2015, Stöckl et al. 2017). In brief, the tracking performance can be described by two components: gain and phase (Farina et al. 1994, Sponberg et al. 2015). The gain is the ratio of the amplitude of the hawkmoth’s movement at the frequency relative to the flower’s movement and would be 1 for perfect tracking. The phase is the amount that the hawkmoth leads or lags the flower movement measured in cycles of oscillations (degrees) and would be 0 for perfect tracking. We used the tracking error ε metric (Roth et al. 2011, Sponberg et al. 2015), which incorporates effects of both gain and phase to quantify the tracking performance of our hawkmoths (Fig.2 –S2). It is calculated as the distance between the moth’s response *H(s)* and the ideal tracking conditions (gain=1, phase lag=0) in the complex plane, where *s* is the Laplace frequency variable:

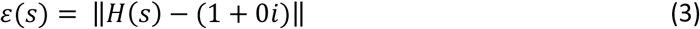

A tracking error of 0, comprised of a gain of 1 and a phase lag of 0, denotes perfect tracking, while at a tracking error larger than 1, the hawkmoths would produce better tracking results when remaining stationary. Because the tracking error metric is represented in the complex plane, we calculated the average of individual tracking errors and their confidence intervals by averaging data in the complex plane to avoid artefacts. These could arise from separating gain and phase components when transforming them and averaging in the non-complex plane (see (Stöckl et al. 2017) for discussion). 95% confidence intervals for gain and phase were calculated in the complex plane as in (Stöckl et al. 2017).

The individual frequency that compose the analyses are not independent measures and instead represent part of dynamic systems characterization of the moth’s response to the flower. Statistical tests are still being developed to compare frequency responses, because it is not clear how to combine data across frequencies. Lacking these, we took the approach of prior studies, which compares the 95% confidence intervals in the gain and phase plots and statistically compares the tracking error and other measures across specific frequency bands. To compare the tracking error across conditions, we pooled the data into two frequency ranges: high and low frequencies, as determined by the frequency range of natural flower movement, which concentrates 95% of total power in frequencies up to 1.7 Hz (as used for other hawkmoth species: (Sponberg et al. 2015)). We used this frequency as the limit for our low frequency category (Fig. 6D), and frequencies higher than 1.7 Hz and up to and including 8.9 Hz into the high frequency range (Fig. 6E). We chose 8.9 Hz as the cut-off for comparability with previous studies on *Macroglossum stellatarum* (Stöckl et al. 2017, Dahake et al. 2018).

### Wing kinematic analysis

To extract the wing beat frequency, as well as the amplitude of wing movement, we analysed the movement of the left wing tip for the first 100 frames of flower tracking. We thereby ensured that we analysed the same stimulus window for all moths to avoid biasing our results by analysing different flight manoeuvres caused by different sections of the stimulus. From these measurements, we extracted the wing beat frequency as the peak in the Fourier transformed wing tip position (Fig. 2 – S1A,B). Because we filmed at a frame rate of 100 fps, the animal’s wing beat frequency was higher than the Nyquist sampling limit, which we corrected for by subtracting the measured frequency from 100 Hz to obtain the actual wing beat frequency. We confirmed this wing beat frequency by filming 3 animals during stationary hovering at 600 fps (wing beat frequencies of these animals: 86.4, 78.4, 77.2 Hz, Fig. 2 – S1C,D). We could not film the tracking videos at 600 fps for all experiments because the memory limits of the camera did not allow sufficiently long videos for the frequency domain analysis.

We used Fiji to extract a horizontal projection of the wing stroke amplitude (Schindelin et al. 2012), measured as the angle between the two most extreme wing positions for each consecutive 10 wing beat sequence over an average of 100 wing beats in total. This resulted in 10 wing beat amplitude measurements per animal, which were then averaged to obtain the wing beat amplitude of the animal (Fig. 2 – S1E,F). This method ensured that we selected the maximum range of wing positions despite the frame rate undersampling of the wing beat frequency. We confirmed the accuracy of our amplitude analysis by comparing the results to the wing beat angles obtained in the brief, 600 fps control videos. With our dorsal camera view, we could not measure the wing stroke amplitude along the wing stroke plane, but measured a horizontal projection of it. Thus, changes in wing stroke angle might appear as changes in the projection amplitude we measured, as the actual wing stroke also extends into the vertical axis (Willmott & Ellington 1997). However, based on unpublished measurements of the wing stroke plane relative to the horizontal plane of *M. stellatarum*, which averages 30.21 ± 3.8 ° IQR, the projection captures the majority of wing stroke amplitude change, and the method is consistent with approaches in other insect studies.

### Aerodynamic model

We used established models of hovering flight (Ellington 1984a, Ellington 1984b, Fernández et al. 2017) to estimate the effects of wing damage on the lift force and mechanical power required for flapping the wings. We modelled the lift production and required mechanical power during flapping hovering for each individual. Following the methods of (Fernández et al. 2017) and (Ellington 1984b) lift production was modelled as:

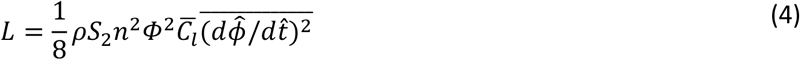

where *ρ* is air density, *S*_2_ is the second moment of wing area, *n* is the wing beat frequency, *Φ* is wing beat amplitude, 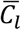 is the average coefficient of lift, and 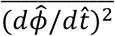 is the average square of the non-dimensional angular velocity. The required mechanical power was modelled as:

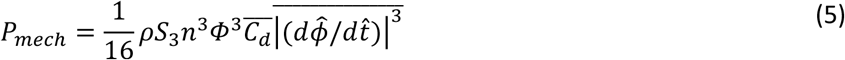

where *S*_3_ is the third moment of wing area, 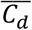 is the average coefficient of drag, and 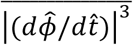 is the average of the absolute value of the cube of the non-dimensional angular velocity. Numerical values of 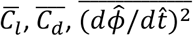, and 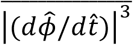 were obtained from (Fernández et al. 2017) and equal to 1.4, 1.6, 19.74, and 105.29, respectively.

Since in these equations, the wing stroke amplitude *Φ* denotes the angular amplitude of the wing stroke in the wing stroke plane, rather than in the horizontal projection as we measured in our setup, and the transformation of the wing stroke angle from projection to stroke plane is not a linear one, we transformed the projected wing stroke angles into the wing stroke plane using the following transformation,

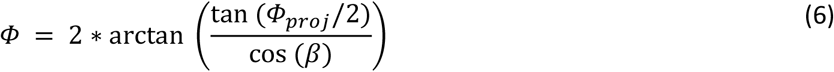

where *β* denotes the angle between the horizontal plane and the wing stroke plane, and *Φ_proj_* is the wing stroke amplitude measured in the horizontal projection. To calculate wing stroke amplitudes back to the horizontal projection for better comparison with our measured results, we used the relationships in equation 6.

To further investigate the relative roles of modulating stroke amplitude and frequency for aerodynamic compensation, we calculated the necessary modulation required in each variable to compensate force production if the other variable was fixed at the control average. In other words, how much greater would the increase in wing beat frequency need to be if stroke amplitude was not modulated (and vice versa). To do this, the lift force was calculated for each individual in the *artificial* damage and *natural* damage group. Using the calculated lift, the equation was rearranged, the mean wing beat frequency from the control group was used to replace the wing beat frequency of that individual, and the resultant magnitude of the wing stroke amplitude was calculated. This procedure was then repeated while using the wing stroke amplitude of the mean control group to determine the wing beat frequency necessary to compensate lift for that individual.

To obtain the required second and third moments of wing area, wing morphology from the photographs of each moth was digitized using the StereoMorph package (version 1.6.2) (Olsen & Westneat 2015) in R (version 3.4.2; The R Foundation for Statistical Computing). The rostral and caudal bases of the left and right forewing were digitized and a series of third order Bézier curves were used to outline each forewing. The curves of each wing were then resampled using the StereoMorph package to generate 50 evenly spaced points (semilandmarks) around the perimeter of each forewing.

The digital shape outputs of the left and right forewing from each moth were further analysed in Matlab (version R2018b – 9.5.0.944444). First, each forewing was rotated so its long axis was perpendicular to the long axis of the body. Wing length, *R*, was measured as the distance between the minimum and maximum value of the wing outline. Wing area was calculated using the ‘polyarea’ function in Matlab. For each wing, the second and third moments of area were calculated following (Ellington 1984a).

## Results

We investigated the effects of wing damage on the hawkmoth *M. stellatarum* in three treatments: a *control* group with intact wings, a group with *“natural”* wing damage that occurred in our flight cages, and an *artificial* damaged group, in which we trimmed the distal forewings to the maximum extent with which the animals were still able to fly (Fig. 1D). The *natural* damage group contained individuals with a wide range of wing damage, thus resulting in the widest spread of forewing area and forewing length of the three treatments (Fig.1–S2B,C, Table S1). We normalised forewing length and forewing area relative to the expected length and size given an individual’s body length, using the allometric scaling relationship between the body length and wing size of intact animals. Relative forewing length and area differed significantly between the *damage* treatments and the *control* group, but not within *artificial* and *natural* damage (Fig. 1E,F, Tables S1 and 2): animals in the *natural* damage group had less than 84.4 ± 14.8 % of the wing length and 78.4 ± 8.5 % of the wing area of an intact animal, while the artificial group had 89.7 ± 5.5 % of wing length and 81.0 ± 7.2 % of wing area. Here and following, spread is reported as interquartile range if not indicated otherwise.

### Wing beat kinematics during hovering flight

To analyse their wing beat kinematics, we filmed individual hawkmoths dorsally as they were hovering at the artificial flower (Fig. 1E,F). The wing beat frequency of the *artificial* damage group was significantly increased compared to the *control* group and *natural* damage group (Fig. 2A, Table S1 and 2), while there was no significant different between the *natural* damage and *control* groups. Similarly, the horizontal projection of the wing stroke amplitude, measured as the angle between the maximum forward and backward extent of the forewing edge in the dorsal camera view, was significantly increased in the two damage groups compared to the *control* (Fig. 2B). Moreover, it was significantly higher in the *artificial* group than the *natural* damage group. In combination, the increased wing beat frequency and amplitude in the damaged groups resulted in a significantly higher wing beat velocity in both groups compared to the control (Fig. 2C).

Since the relative wing length and wing area did not just vary across, but also within treatment groups (Fig. 1D,E), we tested for correlations between wing anatomy and wing beat parameters across treatment groups. We observed a significant linear correlation between relative forewing length and wing beat frequency, wing beat amplitude and wing beat velocity (Fig. 3A-C). Similarly, all wing beat parameters correlated significantly with the relative wing area (Fig. 3D-F), though the variation explained was slightly lower.

**Fig. 3.**
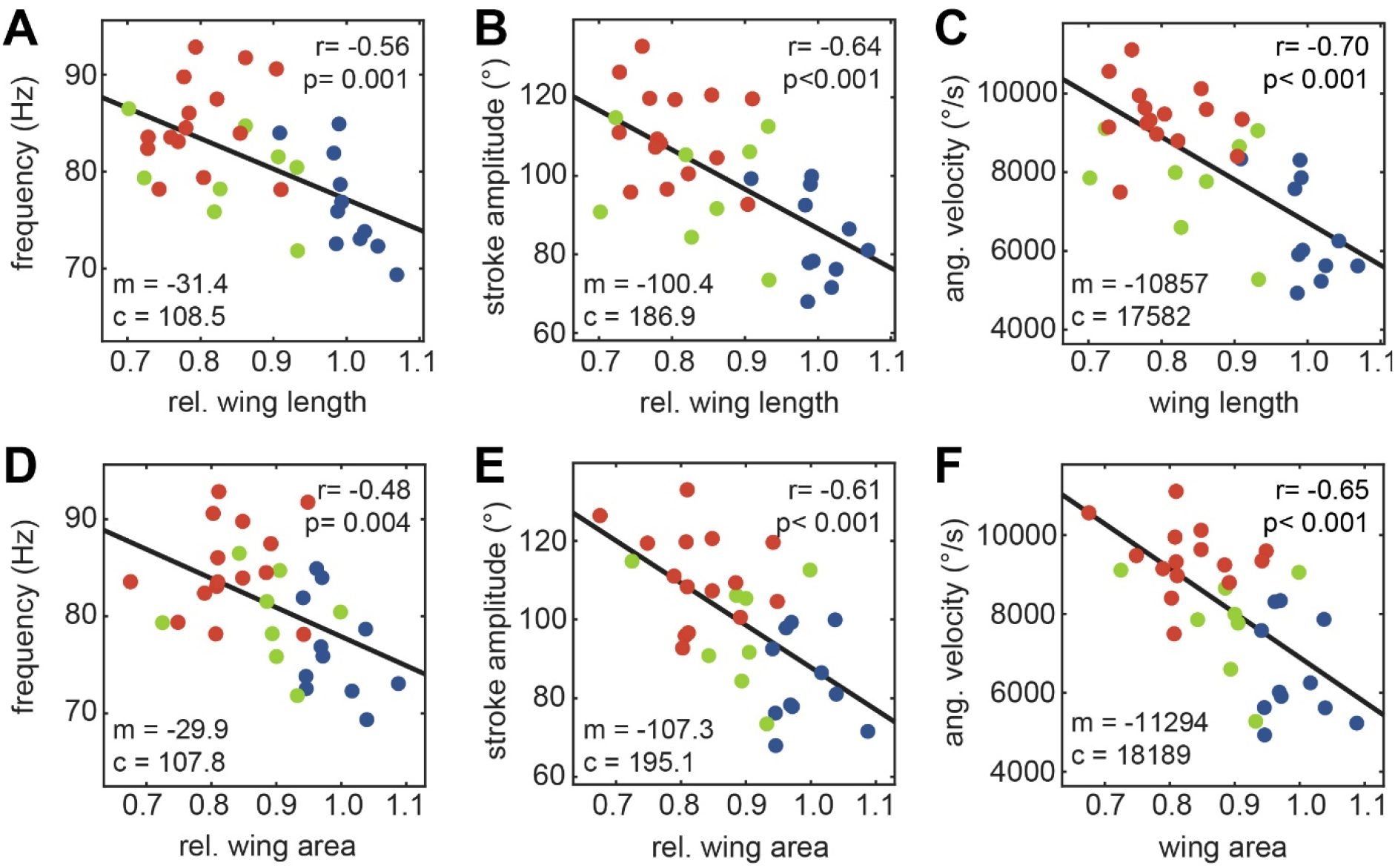
Linear relationship between wing beat kinematics and wing anatomy. Ordinary least square regression of wing beat frequency (**A,D**), horizontal projection of wing stroke amplitude (**B,E**) and wing beat velocity (**C,F**) with the relative forewing length (**A-C**) and area (**D-F**). The strength of the linear correlation is given by *r*, and the statistical significance of the Pearson correlation coefficient by *p*. The model for the linear fit with slope m and intercept *c* is given in each panel.

We also observed a change in the body pitch angle between treatments, measured as the distance between the thorax and the distal tip of the abdomen in the dorsal camera view. Hawkmoths in the *artificial* showed a steeper body pitch angle than animals in the *control* group (Fig. 2-S2A). There was a linear correlation between body pitch angle with relative wing length (Fig. 2-S2B), and relative wing area (Fig. 2-S2C).

### The effect of wing beat kinematics on lift and mechanical power

To test whether the increase in wing beat amplitude and frequency in the *damage* groups could compensate for the loss in lift force due to the reduced wing area, we calculated the effects of these parameters on the lift force and mechanical power required for flapping the wings (see *Methods - aerodynamic model*). There was no significant difference in the estimated lift force across treatments (Fig. 4A, Table S4), indicating that the changes in wing beat amplitude and frequency observed in the *damage* treatment groups were sufficient to compensate for the loss in lift force due to the reduction in wing area. There was no significant difference between treatments in the mechanical power based on the measured wing area and wing beat kinematics (Fig. 4B, Table S4).

**Fig. 4.**
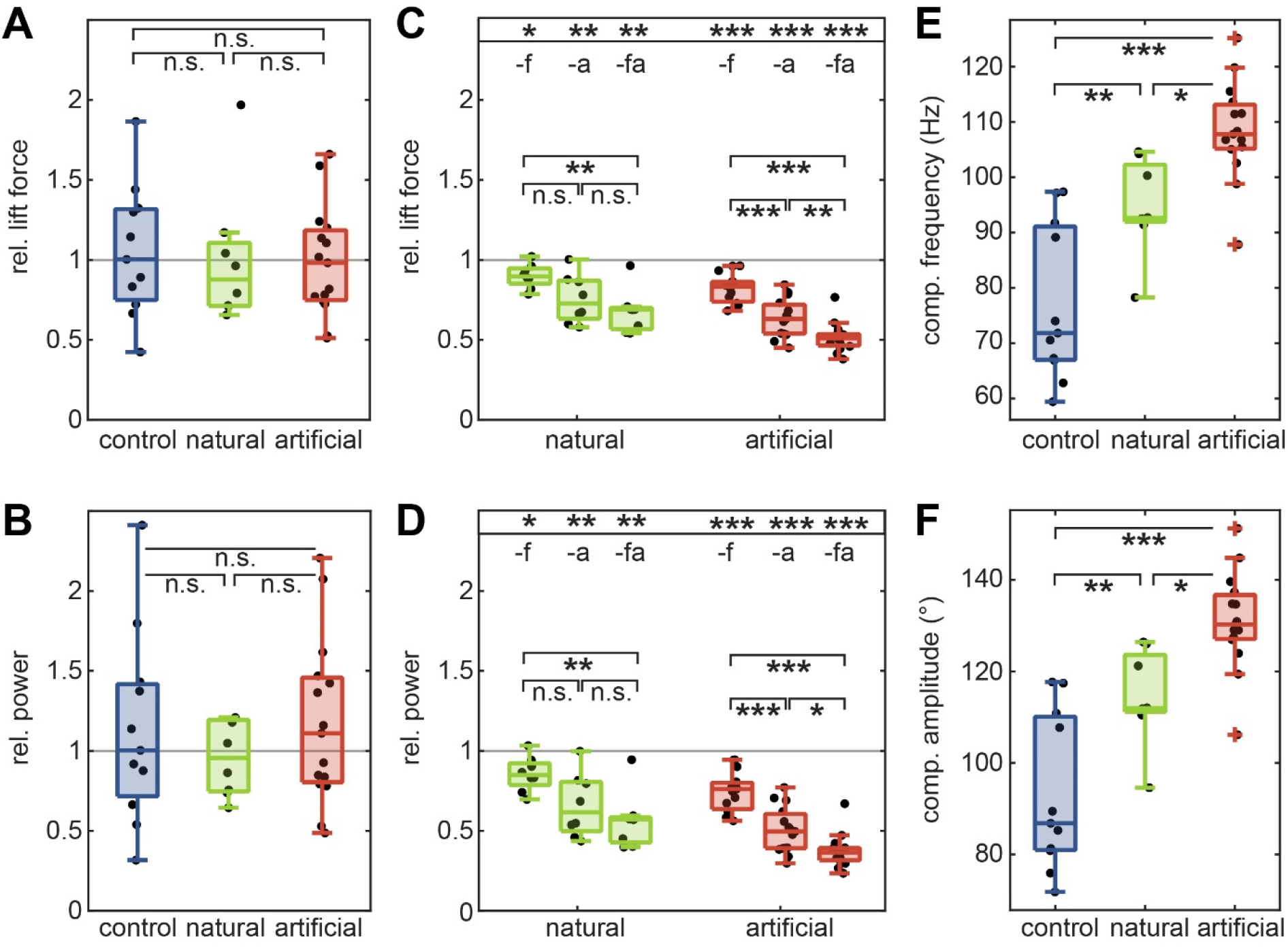
Estimated lift generated, and mechanical power required for flapping. We calculated the lift force generated during hovering (**A**) for the three treatment groups, as well as the mechanical power (**B**) required for flapping, based on their wing shape, wing beat frequency and wing stroke amplitude in the wing stroke plane. Results were normalised relative to the median of the control group. We compared the estimated lift force (**C**) and mechanical power (**D**) within each treatment for three scenarios: with the measured wing beat amplitude (transformed to the stroke plane) and the median wing beat frequency of the control group (*-f*), with the measured frequency and median amplitude of the control group (*-a*) and with both median frequency and amplitude of the control (*-fa*). For each treatment, the resulting lift force and power predictions were normalised by the treatments predictions based on the measured parameters. Using the fixed median wing stroke amplitude of the control group, we calculated the wing beat frequency required to generate the same lift as with both the measured amplitude and frequency (**E**), and vice versa for the wing stroke amplitude (**F**). Statistical differences between groups are indicated as: *** p < 0.001, ** p < 0.01, * p < 0.05, (Kruskall-Wallis test performed after testing the linearity of residuals, see Tables S3, S4, S6). In **C** and **D**, a Wilcoxon signed rank test was used to compare the median of each group with 1: *** p < 0.001, ** p < 0.01, * p < 0.05, n.s. p > 0.05, see Table S6.

To determine if wing beat frequency or wing stroke amplitude contributed more strongly to lift compensation, we replaced the measured wing beat parameters by the median frequency and amplitude of the *control* group. Without any compensation, the loss of lift force was substantial: the median lift force in the *natural damage* group decreased significantly to 66.4 ± 15.2% of the lift force generated with their measured wing beat parameters, and in the *artificial damage* group to 50.0 ± 8.5% (Fig. 4C, Table S5). Wing beat amplitude contributed significantly more strongly to lift compensation than wing beat frequency in the *artificial* damage groups (Fig. 4C, Table S5), as the reduction upon fixing wing beat amplitude in the model was significantly larger than upon fixing wing beat frequency. However, both wing beat parameters contributed significantly to lift compensation, as the resulting lift forces with either one fixed were significantly reduced. This was also the case for the *natural* damage treatment, though the lift force upon fixing wing beat amplitude was only marginally significantly different from fixing wing beat frequency (p=0.052, Table S5, Fig. 4C). Because of the proportional scaling of the lift force and mechanical power equations (see Methods, equations 4 and 5), the effects of fixing wing beat amplitude and frequency on the resulting mechanical power were similar to those on the lift force generation (Fig. 4D).

To investigate the contribution of the two wing beat variables further, we calculated the modulation required in each variable to compensate lift force production if the other variable was fixed at the *control* average. In other words, we calculated how much greater the increase in wing beat frequency would need to be if stroke amplitude was not modulated (and vice versa). It is important to note that the stroke amplitude in our model was assumed in the wing stroke plane, rather than in the horizontal projection as measured in our experiments. We therefore transferred the projected amplitudes to the stroke plane using an average stroke plane angle of 30.21 °. If stroke amplitude was fixed, the average wing beat frequency required to compensate lift force production was 92.6 ± 10.3 Hz for the *natural*, and 107.8 ± 8.0 Hz for the *artificial* damage group, both significantly increased compared to the control (Fig. 4E, Table S7). When wing beat frequency was fixed, the average wing beat amplitude required to compensate lift force production was 111.9 ± 12.5 ° for *natural* damaged wings and 130.3 ± 9.6 ° for *artificial* damaged wings.

### Tracking performance

To quantify the consequences of wing damage for manoeuvrability during hovering flight, we used an artificial flower stimulus simultaneously moving at different temporal frequencies (Fig. 1B). To quantify the hawkmoth’s flower tracking performance, we calculated the absolute Euclidian distance between hawkmoth and flower for the length of the stimulus. There was no significant difference in average displacement across treatments (Fig. 5A, Table S8), nor in peak displacement (Fig. 5B, Table S8). Moreover, hawkmoth flight pathlengths, relative to the pathlength of the flower, did not differ between treatments (Fig. 5C, Table S8), neither was there a significant linear correlation of either of these measures with the wing length of the animals (Fig. 5D-F).

**Fig. 5.**
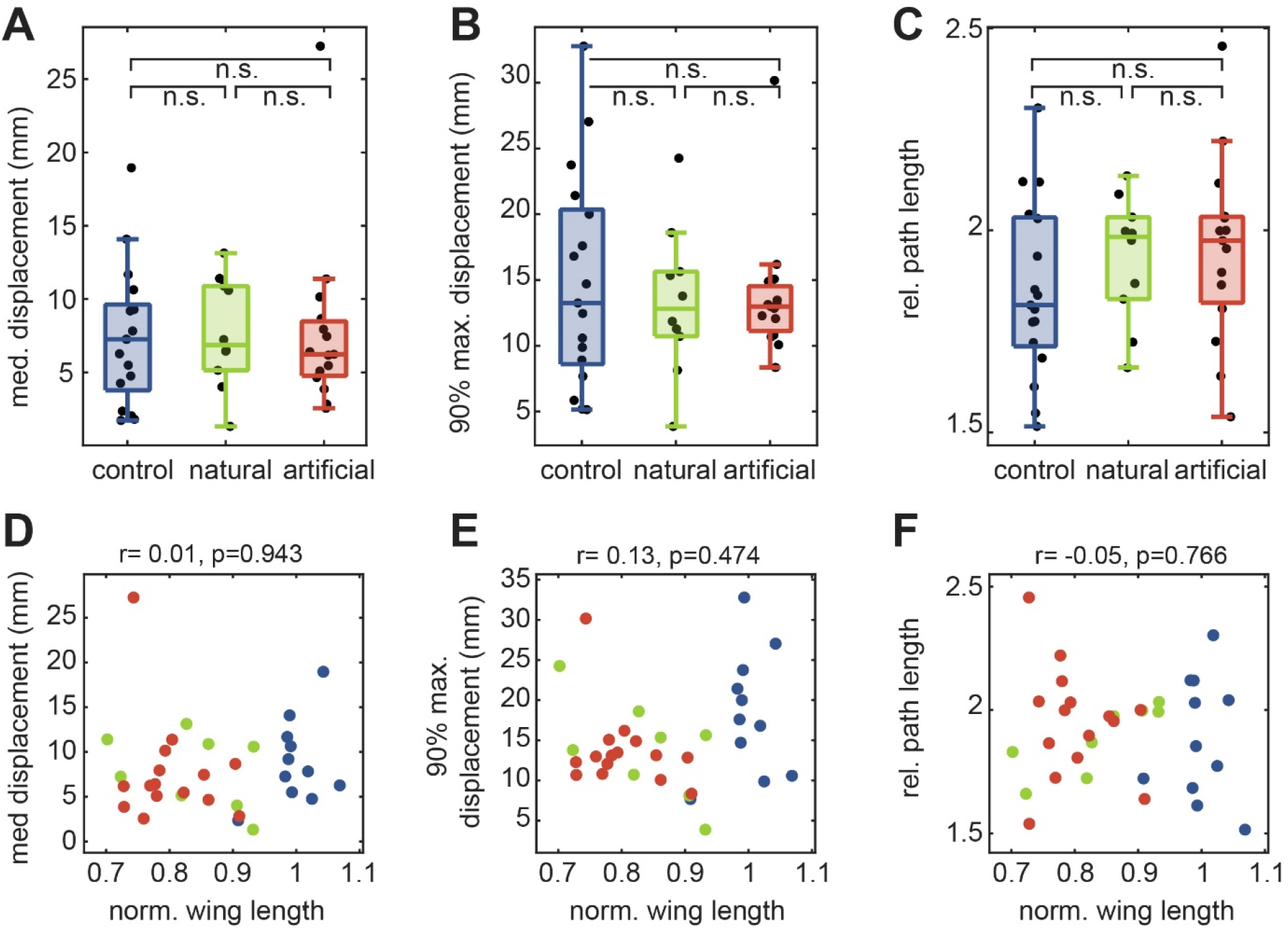
Flower tracking performance of the treatment groups. Median displacement (**A**), 90% maximum displacement (**B**), measured at the Euclidian distance between the hawkmoth’s thorax and flower nectary, and flight path length (**C**) of the control, natural and artificial wing damage groups. Statistical differences between groups are indicated as: n.s. p > 0.05 (ANOVA with Tukey’s HSD corrected post-hoc test performed after confirming the linearity of residuals, Table S8). **D-F** Linear correlation between normalised wing length median displacement (**D**), 90% maximum displacement (**E**) and flight path length (**F**). R indicates the strength of the linear correlation, and p the statistical significance of the Pearson correlation coefficient.

To analyse the flower tracking performance for the combined frequency response of the moth in detail, we extracted the gain and phase of the hawkmoth’s response (Fig. 6A,B). Like in previous studies on this species (Stöckl et al. 2017, Dahake et al. 2018), we observed tracking responses with distinct gain and phase characteristics: a gain overshoot between 2 and 4 Hz, as well as a plateau of the gain between 6 and 11 Hz. This secondary plateau had a higher gain in the *artificial* damage group compared to the *control* and *natural* damage group with non-overlapping confidence intervals (Fig. 6A).

**Fig. 6.**
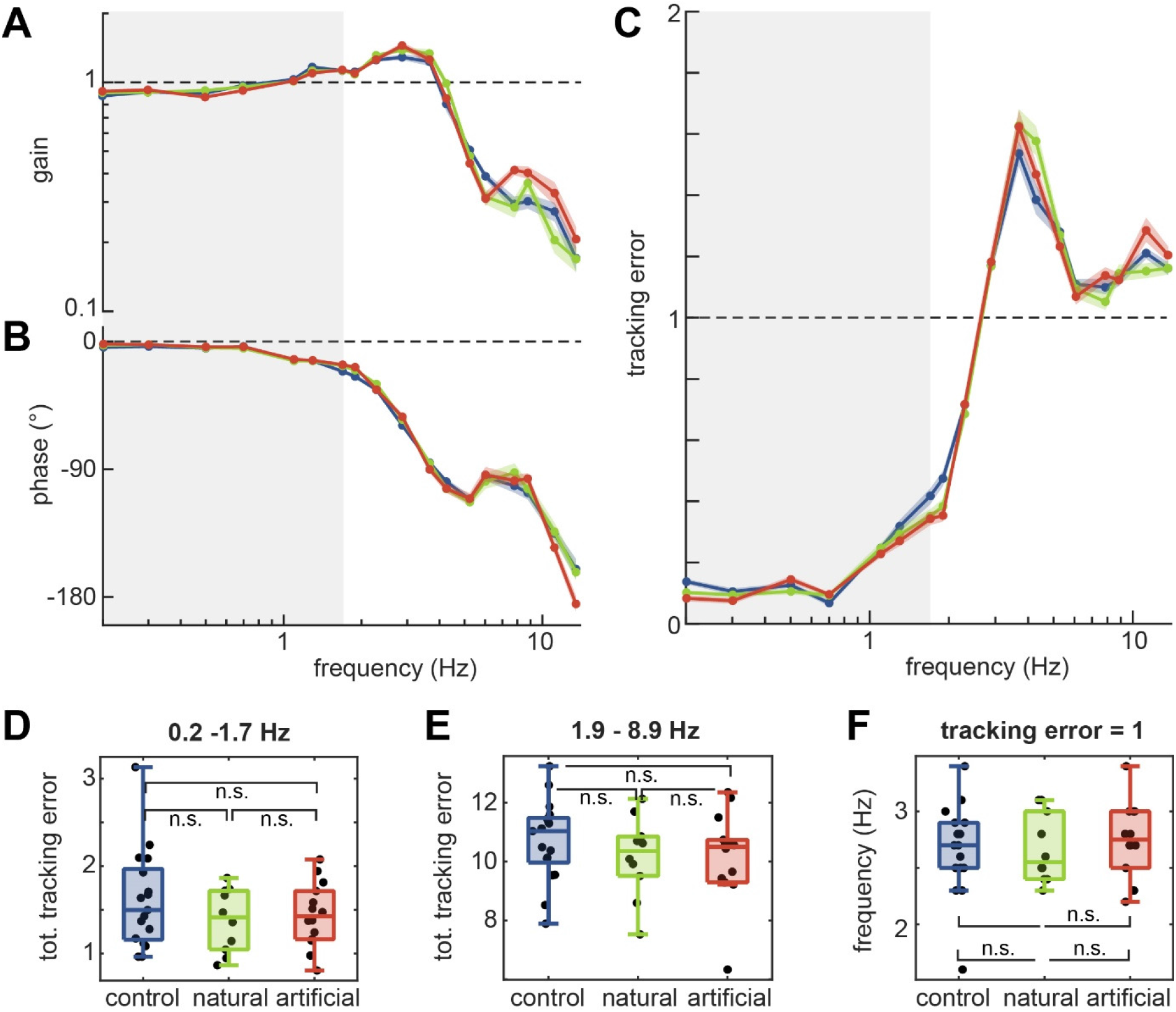
Flower tracking performance of the treatment groups at different temporal frequencies. Gain (**A**), phase (**B**) and tracking error (**C**) of the control, natural and artificial wing damage groups. Curves show the mean and 95% confidence intervals of the mean computed in the complex plane. The total tracking error for each group over a low (**D**) and a high (**E**) temporal frequency range. **F** The temporal frequency at which the tracking error approached 1 for each treatment group. Statistical analysis was performed using ANOVA with Tukey’s HSD corrected post-hoc test (**D,F**) and a Kruskal-Wallis test (**E**) after testing the linearity of the residuals. ** * p < 0.01, * p < 0.05, n.s. p > 0.05. See Table S9. The grey areas represent the lower temporal range of tracking frequencies analysed in **A.**

From the combination of gain and phase we calculated a tracking error metric (see *Methods*), to quantitatively compare the flower tracking performance of the different treatment groups (Fig. 6C). The average tracking error of the different treatment groups was very similar: it was low for frequencies below 1 Hz, and then rose to 1 at about 2.5 Hz. Correspondingly, the total tracking error in the lower frequency range, in which 95% of the power of flower movements is concentrated (Sponberg 2015), did not differ significantly between treatment groups (Fig. 6D, Table S9), neither did it differ in the high frequency range 1.9 to 8.9 Hz (Fig. 6E, Table S9). The total tracking error integrated over all temporal frequencies did not correlate with the relative forewing length of forewing area (Fig. 6 – S1A,B) and neither with absolute animal size (Fig. 6 – S1C). An error of 1, above which the animal would do equally well by staying stationary, was reached at the same frequency for all treatment groups, indicating that there was no significant difference in the temporal response of the different groups (Fig. 6F, Table S9).

## Discussion

### Hummingbird hawkmoths use a combination of frequency and amplitude to compensate for wing damage

In our study on the effects of wing damage on flight performance in the hawkmoth *Macroglossum stellatarum* we observed significant effects of wing damage on the flight kinematics. Similar to other studies previously performed on the hawkmoth *M. sexta* (Fernández et al. 2012, Fernández et al. 2017), as well as on butterflies (Kingsolver 1999) and bumblebees (Hedenström et al. 2001, Haas & Cartar 2008), hummingbird hawkmoths increased their wing beat frequency when their wings were damaged (Fig. 2A). The wing beat frequency was increased proportionally to the amount of forewing area and length lost (Fig. 3). The increase in wing beat frequency in wing-damaged hummingbird hawkmoths was larger than in *M. sexta* (10% compared to less than 5%, (Fernández et al. 2017)), and larger in relative terms than the changes in wing beat frequency observed in bumblebees upon comparable extent of wing damage, though similar in absolute extent (Hedenström et al. 2001, Haas & Cartar 2008). Thus, our findings show that wing beat frequency can also be flexibly adjusted in insects with synchronous flight muscles, to a similar extent as in insects with asynchronous flight muscles.

Interestingly, we also observed a significant and substantial increase in wing beat amplitude in hawkmoths with damaged wings (Fig. 2B). A similar increase in wing beat amplitude has not been observed in any of the species so far studied – though there was a marginally significant increase in wing beat amplitude in *Manduca sexta* (Fernández et al. 2017). The increase in *M. stellatarum*, however, was much larger than in *M. sexta*, and reached about 30%, while in *M. sexta* it was only 5%. Together, our model results indicate that in *M. stellatarum*, the adjustments in wing beat kinematics are sufficient to compensate for the loss in lift force in the *damage* treatments, without a significant increase in mechanical power required to move the wings (Fig. 4).

Why, though, did hummingbird hawkmoths show a much more pronounced change in wing beat kinematics to compensate for a loss of wing area than their larger relative *M. sexta?* A striking difference in flight kinematics between *M. sexta* and *M. stellatarum* is their average wing beat frequency, and their overall difference in size (*M. sexta* has more than twice the wingspan of *M. stellatarum* and on average weighs 5 times as much (Henningsson & Bomphrey 2013)). However, the general shape of the wings across hawkmoths is very similar, suggesting similar hovering kinematics. Their wing loading and span efficiency have been shown to be similar (Henningsson & Bomphrey 2013). Yet, this assessment was based on experiments performed on tethered individuals in a wind tunnel, so the results might differ from those for freely hovering individuals. Indeed, the wing beat frequency of freely hovering *M. stellatarum* is almost twice as high as that measured in tethered animals (76 Hz in Fig. 2 vs 48 Hz, (Henningsson & Bomphrey 2013). Thus, further kinematic studies investigating hovering flight in diverse hawkmoths will be necessary to understand the striking differences in wing beat kinematics following wing damage in these two hawkmoths species.

### The high wing beat frequency of M. stellatarum might require a compensation strategy via wing beat amplitude

Why did we observe a much greater contribution of wing beat amplitude in lift force compensation than previously observed in other insects? Our modelling gives some indication for why *M. stellatarum* might use this strategy, rather than compensate almost entirely by adjusting wing beat frequency, as most other insects do. Based on our calculations, in order to compensate the reduction in lift force due to wing damage entirely by adjusting wing beat frequency, an average frequency of 93 Hz for the *natural* and 108 Hz for the *artificial* group would be required (Fig. 4E). However, such wing beat frequencies might exceed the range of synchronous flight muscles (Dudley 2000, Syme 2002), the muscle type possessed by hawkmoths. Indeed, none of the hawkmoths in our study (in any treatment group) reached even the more moderate wing beat frequency increase of 94 Hz required for the *natural* damage group (Fig. 2A). Since the wing beat frequency of intact *M. stellatarum* is already quite close to the upper limit for synchronous flight muscles, compensating lift force production entirely by wing beat frequency might not be an option for this hawkmoth species, and therefore a reason why we observed a larger contribution of wing beat amplitude to lift force compensation (Fig. 4C). The significant contribution of wing beat frequency, nevertheless, suggests that wing beat amplitude alone might not be sufficient to compensate the reduction in lift force upon wing damage. It is therefore likely that the required increase in wing beat amplitude (especially in the *artificial* damage case) might be outside biomechanical limits at the relatively high wing beat frequencies employed by *M. stellatarum*.

A potential beneficial side-effect of this compensation strategy, which relies heavily on increases in wing beat amplitude rather than frequency, is a reduction in the inertial power required to move the wings. While the aerodynamic power (Fig. 4B) scales similarly for changes in wing beat frequency and amplitude, inertial power scales with the square of stroke amplitude and the cube of wing beat frequency (Willmott & Ellington 1997). Inertial power is often not considered as a cost, because it is assumed that minimal power is required for the inertial acceleration of the wings due to energy storage and return. However, it has been shown that the inertial power, while significantly reduced, is not perfectly compensated by elastic elements (Gau et al. 2019). Thus, the amplitude-based strategy of *M. stellatarum* should be beneficial from an inertial power point of view, and equally good as far as aerodynamic power is concerned. This raises the question, however, why all other insect species investigated so far showed a frequency-based compensation strategy. One important aspect speaking against an amplitude-based compensation strategy, in particular for insects with asynchronous flight muscles, is that modulations of wing beat amplitude are required for steering. Thus, increasing the average wing beat amplitude proportionally leaves less room for amplitude changes, which might therefore affect manoeuvrability in these insects. It will therefore be interesting to investigate in the future whether insects with synchronous and asynchronous flight muscles apply different strategies when compensating for wing damage, and what role other factors, such as body and wing size, might play.

### Flower tracking manoeuvrability is not compromised by natural or artificial wing damage

While the effects of wing damage on flight kinematics show some common trends across the different species of insects studied previously, it is less well understood what effects wing damage has on the manoeuvrability of fast flying insects, and thus ultimately on their foraging and predator avoidance success. In our setup, we studied the effects of wing damage on the manoeuvrability of hawkmoths in a task which is very similar to their natural foraging paradigm. Moreover, we could test the effect of wing damage quantitatively over a range of temporal frequencies, and thus could see whether wing damage affected particular temporal aspects of the hawkmoth’s flower tracking ability, as does alteration of their sensory input (Sponberg et al. 2015, Roth et al. 2016, Stöckl et al. 2017, Dahake et al. 2018). However, we did not find any significant impairment of the tracking performance in wing-damaged hawkmoths (Figs. 6,7), and no significant difference in tracking error within the frequency range in which flowers naturally move ((Sponberg et al. 2015), Fig. 6D), and nor indeed at any frequency across the entire range. We did, however, observe an interesting difference between the three treatment groups in the shape of their tracking response, in particular in the tracking gain: individuals with *artificial* wing damage had higher tracking gains for frequencies ranging from 4.5 to 12 Hz (Fig. 6A). The particular shape of the frequency response (Fig. 6A,B) is a representation of the full sensory to motor dynamics of the hovering moth. These differences could therefore arise either from changes in flight kinematics affecting the mechanics of flight, or from context-dependent changes in neural processing due to damage.

Even though these differences did not affect overall tracking error, they might still reflect functional differences in the control and damaged conditions. Previously, changes in gain within a species were observed upon altered sensory input necessary for flight control, for example when the animals had reduced luminance (Sponberg et al. 2015, Stöckl et al. 2017), conflicting mechanosensory and visual cues (Roth et al. 2016) or were deprived of fast sensory input about the animal’s own movements (Dahake et al. 2018). Without the latter form of feedback, the animals could not track fast movements of the flower, and the gain in the high frequency range decreased compared to the controls. Interestingly, with wing damage, the gain increased in the high frequency range compared to the control group, suggesting that wing damaged animals performed coordinated flight manoeuvres with larger amplitudes within this range. A potential explanation for this observation might be the increased wing beat frequency observed in the wing damage groups. It might allow the animals to perform more accurate flight manoeuvres even at higher frequencies because the necessary adjustments of the hawkmoth’s position could be performed faster. While the effects were relatively small and manifested only outside of the range the animals usually experience, it shows that manipulating the wing anatomy could also be used to artificially change different aspects of wing beat kinematics and study their role in fine-scale flight control.

Overall, our findings suggest that intact wings are not crucial for the precise control of lateral flight manoeuvres, which hawkmoths perform during flower tracking. This is in line with results from foraging butterflies and bumblebees, which likewise showed no significant alterations in flight or foraging performance (Kingsolver 1999, Haas & Cartar 2008). The system identification approach we used on the sum-of-sines stimulus allowed us to extend this conclusion over the full temporal frequency response of the moth’s behaviour. One explanation that might reconcile our results and previous findings from dragonflies, which showed a strong impairment in flight performance upon wing wear (Combes et al. 2010) is the direction in which flight manoeuvres were executed. In our experiments, the insects were conducting horizontal flight manoeuvres, while in dragonflies the vertical acceleration was impaired, and animals often perform vertical manoeuvres during prey capture, which showed a reduction in success upon wing damage (Combes et al. 2010). One might therefore hypothesise that wing damage specifically affects flight manoeuvres that require fast vertical but not horizontal acceleration.

### Conclusion

Taken together, hummingbird hawkmoths compensate for a loss in wing area by increases in wing beat frequency and amplitude, and track moving flowers without a performance impairment. This impressive tolerance to wing damage might be a testament to the immense importance that fast steering has for these animals: not only do they feed exclusively on the wing, and very rarely land on flowers, they also lay their eggs on their hostplant while hovering in front of the plants (Stöckl & Kelber 2019). Moreover, since hummingbird hawkmoths hibernate as adults, resulting in lifespans of several months (Pittaway 1993), optimising their flight abilities to tolerate wing damage might be paramount for the fitness of these insects. Their strategy to compensate for the loss in wing area both by an increase in wing stroke amplitude and wing beat frequency, suggests that multiple kinematics strategies could be utilized to compensate for wing damage in different insect species. It opens up future study directions to better understand which kinematic, aerodynamic and behavioural aspects govern the strategies of insects to compensate for wing damage while retaining optimal manoeuvrability.

## Acknowledgements

We thank Merry and Leigh Foster for help with capturing the parental moths in France, and Marie Dacke for allowing us to use her high-speed cameras. We thank Usama Bin Sikandar for help with analysis.

## Author contributions

Conceptualization: A.S., K.K.; Methodology: A.S., S.S.; Validation: K.K., A.S.; Formal analysis: K.K., A.S., B.A.,nS.S.; Investigation: K.K.; Resources: S.S., A.S., E.W.; Data curation: K.K., A.S.; Writing - original draft: A.S.; Writing - review & editing: A.S., K.K., E.W., S.S., B.A.; Visualization: A.S.; Project administration A.S.; Supervision: A.S., S.S.; Funding acquisition: E.W., S.S., B.A.

## Data availability

Data supporting the presented results is confidentially available at Figshare (https://fiqshare.com/s/73ad11df08567ea76d9d) and will be publicly available upon acceptance of this article.

